# Hybrid TF-IDF and SBERT Feature Engineering with SMOTE-Tomek Resampling for Enhanced Software Requirements Classification

**DOI:** 10.64898/2026.01.13.699413

**Authors:** Hashim Ali, Raja Sarath Kumar Boddu, Umer Tanveer, Aamir Saeed, Yazeed Yasin Ghadi, Masoud Alajmi, Khalfalla Awedat, Hend Khalid Alkahtani

## Abstract

Software requirements classification remains one of the important challenges in requirements engineering. Engineering that affects the smoothness of project success about software development life cycles. in this paper, a novel hybrid solution is being presented that beats the benchmarks set by previous approaches using TF-IDF. The vectorization, coupled with SBERT embeddings, has been further reinforced by application of SMOTE-Tomek resampling. The solution addresses typical limitations detected in Or’s baseline solution, which achieved 76.16% ± 2.58% accuracy of the model using the conventional SMOTE-Tomek pre-processing with logistic regression. Via comprehensive experimentation on the PROMISE dataset containing 969 categorized requirements accross 12 classes, our hybrid feature engineering approach has shown significant improvements: Random Forest achieves. Thus, the character CNN records an accuracy of 74.40% ± 3.00% versus the baseline 60.37% ± 2.76%, while critically, Naive Bayes shows remarkable improvement from complete failure 0.00% to 53.97% ± 5.08% accuracy. The integration of semantic contextual understanding through SBBERT, with statistical term importances via TF-IDF, forms a robust representation space that captures the synthetic pattern along with semantic relations in the texts of requirements. Our methodology uses stratified 10-fold cross-validation with Matthews Correlation Coefficient (MCC). Evaluation, Ensuring reliable performance assessment across imbalanced classes. The results demonstrate that hybrid feature engineering, when combined with appropriate resampling techniques, provides a more effective solution for automated software requirement classification than traditional pre-processing methods alone.

## 1 Introduction

Software Development Life Cycle (SDLC) lies upon Software requirement engineering, Where accurate requirement classification into either Functional or Non Functional affects success of a project, resource allocation, and stakeholder sanctification [36, 43]. The exponential growth in software complexity and the increasing demand for automated requirements processing have elevated the importance of machine learning-based classification systems [38, 39]. Traditional yet manual methods for requirement classification are exhaustive but very time consuming, prone to human errors, and do not scale well for large projects. [44, 45].

Recent advances in NLP and machine learning have opened up new opportunities for the automation of requirement classification. But at the same time, contemporary methods face serious, the challenges include class imbalance in requirement datasets and limited semantic understanding of technical texts. [3, 4]. These challenges have been attributed to meaningless keywords and phrases, inadequate modeling of context relations inherent in the requirement text representations, and poor feature representations.Improvements in performance were noted in the work of Or’s [1] with SMOTE-Tomek preprocessed data, reporting 76.16% ± 2.58% using Logistic Regression on the PROMISE data set. While this represents progress, with an approach that fundamentally relies on classical vectorization techniques that fail to capture the semantic depth in requirement specifications. The advent of transformer-based models, especially Bidirectional Encoder Representations, the appearance of Transformers-more specifically, from transformers (BERT) [11] and its sentence-level variant SBERT [2], has revolutionized text. Representation: This captures contextual relationships and semantic understanding that were previously out of reach using statistical methods. SBERT is designed for sentence-level. It describes the embeddings that address the computational limitations of BERT while preserving semantic richness, essential in requirement classification tasks.

The following research work, thus, presents a new hybrid solution combining TF-IDF vectorization with Improved SBERT embeddings, which use the modified SMOTE-Tomek resampling to overcome state-of-the-art solution limitations. The contributions of our work are three-fold:

(1) A new solution: providing an overall framework that creates features from both importance measures of statistical terms and meaningful relationships of context in requirements, (2) proof that our solution provides better results in comparison to the existing methods for a variety of machine learning models, (3) critical discussion on MCC metrics for requirement evaluation with high level of imbalance in the requirement classification tasks, and (4) real-world insights to implement automatic classification needs in various industrial-grade software development processes.

The experimental assessment on the dataset of PROMISE shows the following strong improvements: Random Forest-accuracy rises from 60.37% to 74.40%, and Naive Bayes transforms from complete failure, 0.00%, into viable performance, 53.97%. These results demonstrate that hybrid feature engineering, if properly implemented with appropriate resampling techniques, provides a more robust and effective solution to automated software requirement classification than traditional approaches.

## 2 Literature Review

### 2.1 Software Requirements Classification

During the past two decades, the process of software requirements classification has transformed from traditional automated approaches based on rules to advanced machine learning approaches. In the past, the initial work of Cleland-Huang et al [36]. provided crucial foundations for the automated classification of non-functional requirements, where modest gains were observesd based on keyword-driven approaches. Afterward, successive studies have incorporated increasingly complex machine learning models to address the complexity of requirements descriptions that exist in subtle details of the requirements.

Canedo and Mendes [3] presented a thorough comparison between text feature extraction methods and machine learning algorithms for requirement classification, setting a benchmark for conventional methods using TF-IDF transformation along with Support Vector Machines (SVM) and Naïve Bayes classifiers. This study demonstrated the key significance of feature engineering for requirement classification tasks, along with the significance of handling class imbalance.

In recent years, attention has been devoted to adopting methodologies in deep learning. The classification of software requirements using deep learning algorithms has been investigated by Khayashi et al. [4], and competitive results were achieved in classifying software requirements for the software datasetPURE using convolutional networks and recurrent networks. Their method required extreme computational powers and lacked interpretability for industrial use cases.

The advent of transformers has also brought in new avenues for classification of requirements. Airlangga et al. [6] studied GAN-BERT models for improved classification of requirements and proved the capability of attention mechanisms in understanding semantically related requirements in natural language requirements. Abbas et al. [8] have used ensemble learning methods by combining different traditional classification models for improved results for different requirements.

### 2.2 Class Imbalance in Requirements Engineering

“Class imbalance” poses a basic problem for the classification of requirements, where functional requirements, as a rule, are much more numerous than other requirements, including security or usability requirements. Rao et al. [9] have considered class imbalance methods in software engineering tasks, clearly illustrating that traditional classification methods are ineffective to properly learn minority classes, producing a class bias mostly leaning to majority classes.

The Synthetic Minority Oversampling Technique (SMOTE) [13] has emerged as a foundational approach for addressing class imbalance by generating synthetic minority class examples. Tomek links [14] complement SMOTE by removing majority class examples that contribute to classification ambiguity. The combination of SMOTE-Tomek has proven to be quite effective for software engineering applications, as shown in Or’s baseline work [1].

Fernández et al. [54] presented an overall description regarding the different SMOTE variations, as well as their efficacies for various domains. The authors have especially highlighted that there is a need for proper hyperparameter tuning for optimal performance, such as some required modification considerations based on domains for improved results. Rana et al. [7] have discussed various oversampling methods specifically for machine learning models, giving a brief insight into choosing a proper oversampling strategy according to various dataset requirements for a classification task.

### 2.3 Transformer Models in Natural Language Processing

The transformer model [12] has made the biggest impact in the natural language processing field, allowing for performance capabilities in text understanding tasks of unprecedented quality. BERT [11] proposed the task of bidirectional context understanding for the masked language modeling task and achieved outstanding performance on a number of natural language processing benchmarks. BERT is computationally heavy; also, it is not the best model for the task of computing the similarity of sentences in the context of requirement classification.

To overcome the mentioned limitations, Sentence BERT (SBERT) [2] was developed by altering the BERT model for obtaining fixed-sized embeddings for sentiment analysis tasks. The method utilizes a siamese network for training with the Natural Language Inference dataset. The approach was found to be more effective for tasks involving the calculation of semantic similarities between two sentences and can be applied for various tasks in different domains.

Currently, some recent works on transformer efficiency include DistilBERT [32], a model that preserves 97% of the performance of the original BERT but with a 40% reduced size, and RoBERTa [31], which focuses on the optimization of the original BERT training procedure to improve robustness. Despite these, SBERT is still the best-performing model for sentence-level requirement classification tasks due to its custom design.

### 2.4 Evaluation Metrics for Imbalanced Classification

Classic evaluation criteria such as accuracy and F1 measure may prove deceptive on imbalanced datasets, which is a very important consideration in the classification of requirements, where some types of requirements inherently have low occurrences. The superiority of the use of the Matthews Correlation Coefficient (MCC) over classic evaluation measures such as accuracy in binary classification tasks was proven by Chicco and Jurman in [5].

The MCC, defined as:

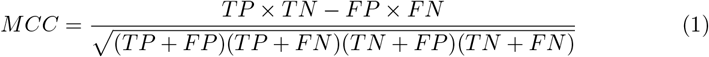

where TP, TN, FP, and FN are true positives, true negatives, false positives, and false negatives respectively, is a balanced measure which takes into account all the facets of the classification. This is different from the F1 measure as it does not ignore the true negatives. This is particularly important when dealing with imbalanced classifications, such as requirement classifications.

## 3 Methodology

It is a general methodology workflow which tends to highlight the systematic approach to a research study which comprises different levels such as data processing, hybrid feature development, resampling techniques, and evaluation.

### 3.1 Dataset Description

The PROMISE (PRedicting Object-Oriented Software Development Effort) is a dataset used experimental base for this study, consisting of 969 software requirements classified into 12 classes, which cover functional as well as non-functional types of requirements. The dataset exhibits significant class imbalance characteristic of real-world requirement collections: Functional requirements (F) constitute 444 instances (45.8%), Security requirements (SE) include 125 instances (12.9%), Usability requirements (US) comprise 85 instances (8.8%), while other categories such as Legal (L) and Portability (PO) contain as few as 15 and 12 instances respectively.

This distribution is imbalanced, reflecting real-world software projects where functional requirements dominate and specialized non-functional requirements occur less often but are critical regarding system quality and user satisfaction. In general, the dataset is a very good case study in order to evaluate the performance of different classification techniques dealing with imbalanced sets of requirements.

### 3.2 Hybrid Feature Engineering Framework

Our hybrid feature engineering method is designed to combine all these in a manner that leverages both the importance of individual terms in a statistical manner utilizing TF-IDF vectorization and understanding in a manner that comprehends the context provided by SBERT embeddings. This complementary method is tailored to address both individual weaknesses in a manner that takes advantage of term frequency patterns and domain-specific importance provided by both methods that is illustrated in Figure 5

**Fig 1.**
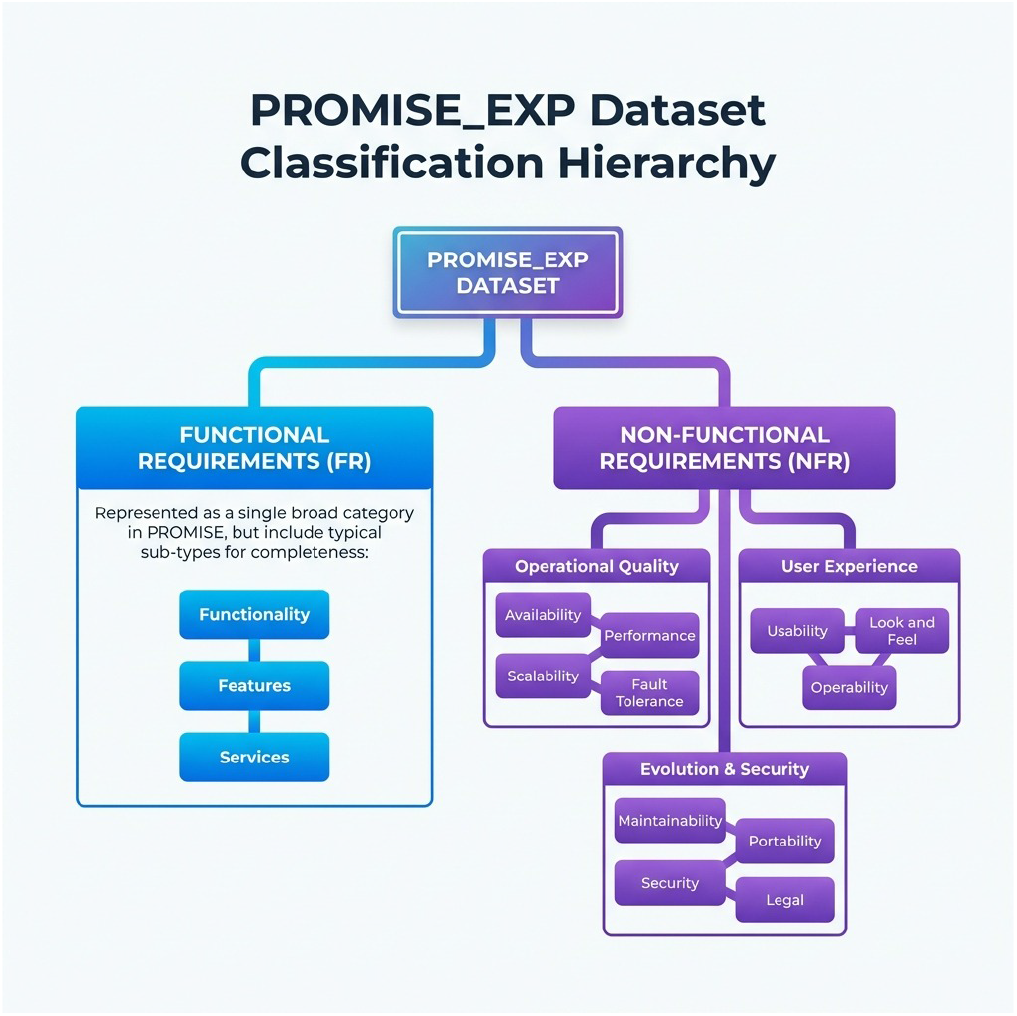
Software requirements taxonomy showing the hierarchical classification of functional and non-functional requirements with their respective subcategories.

**Fig 2.**
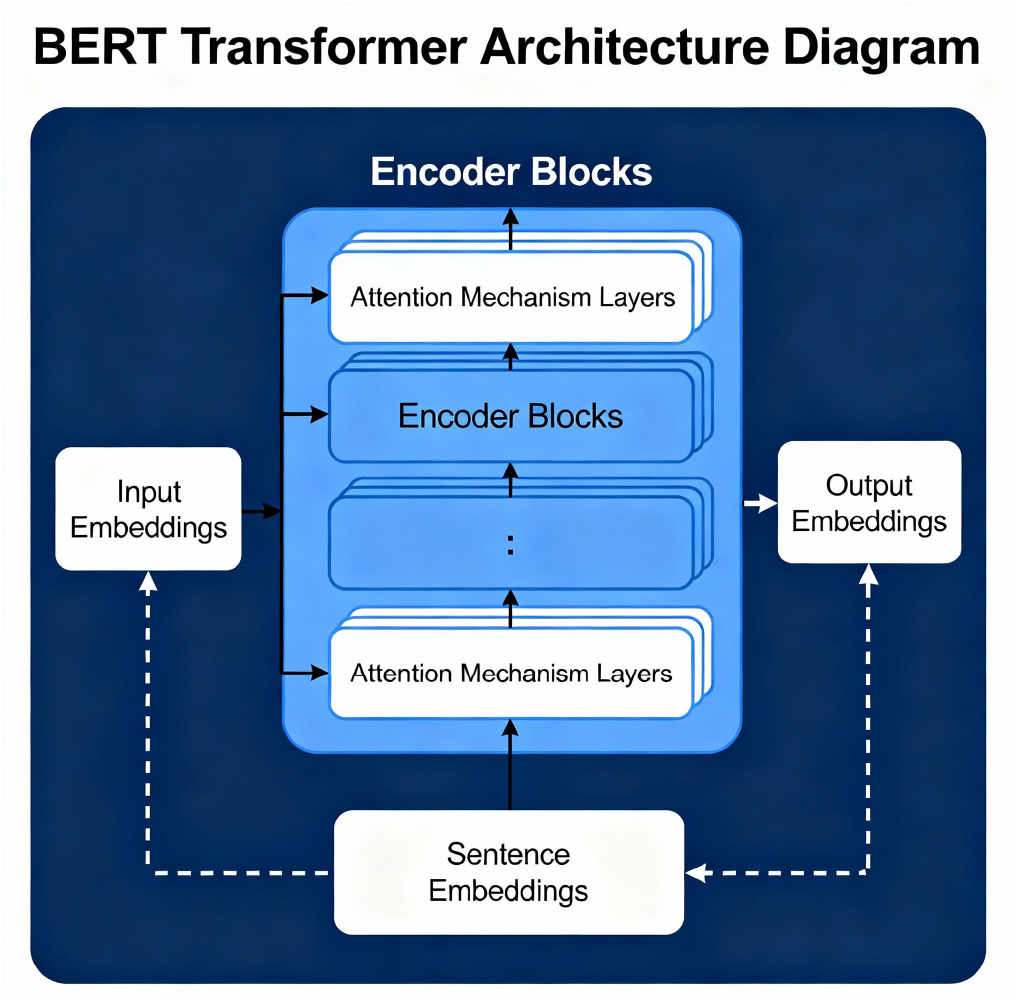
BERT transformer architecture illustrating the multi-layer encoder structure with attention mechanisms that enable bidirectional context understanding for sentence-level embeddings.

**Fig 3.**
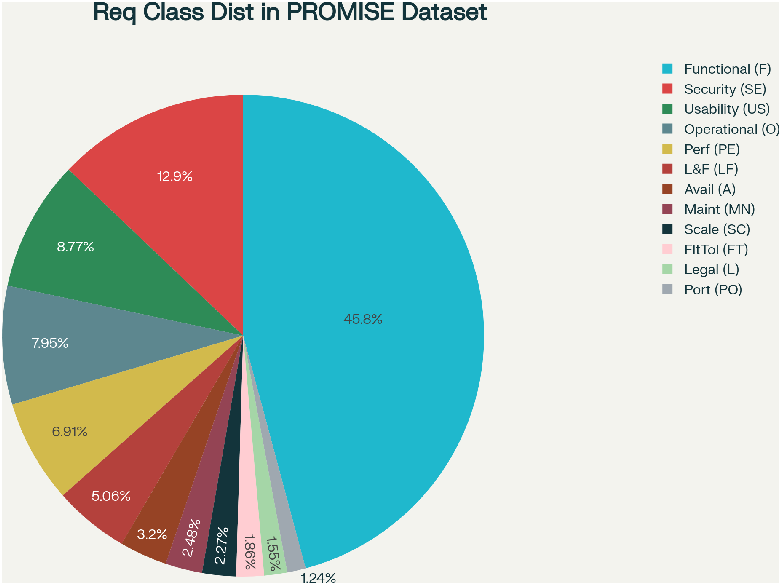
Distribution of requirement classes in the PROMISE dataset, illustrating the significant class imbalance with functional requirements comprising 45.8% of all instances while specialized requirements like legal and portability represent less than 2% each.

**Fig 4.**
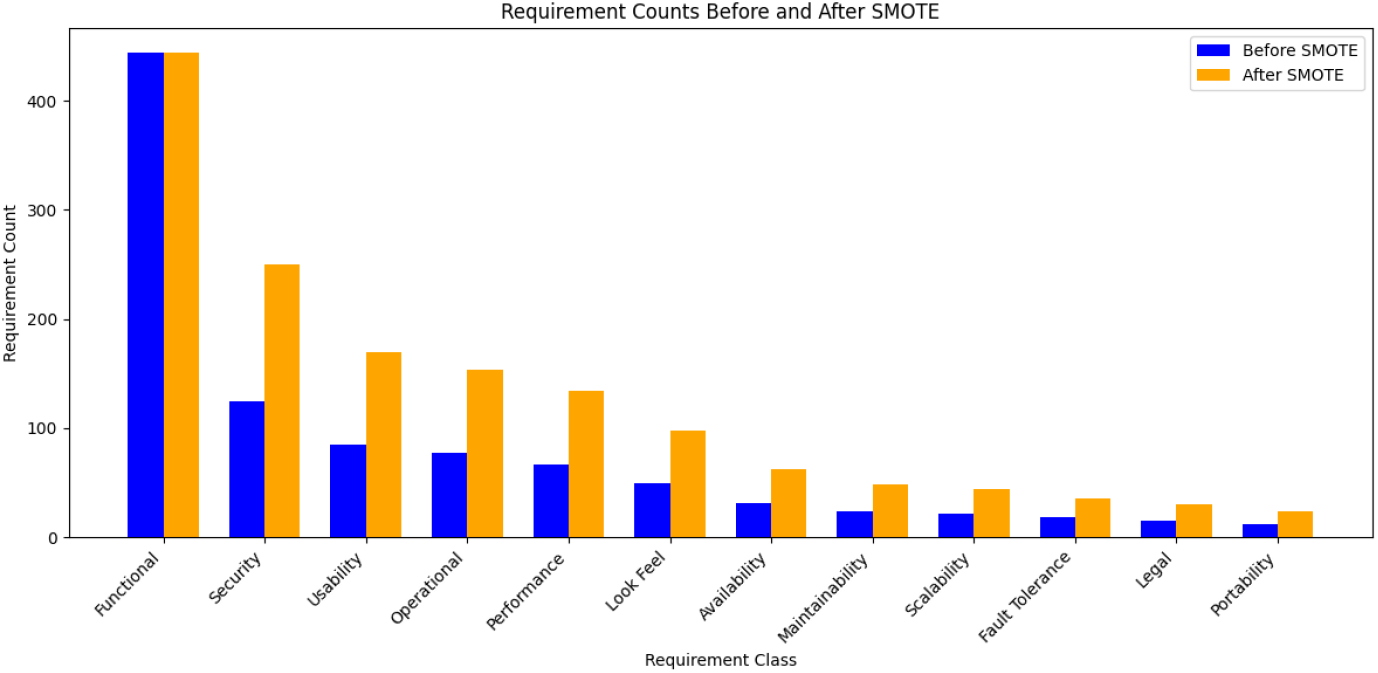
SMOTE oversampling technique illustration showing how synthetic minority class samples are generated through linear interpolation between existing minority instances and their k-nearest neighbors to address class imbalance.

**Fig 5.**
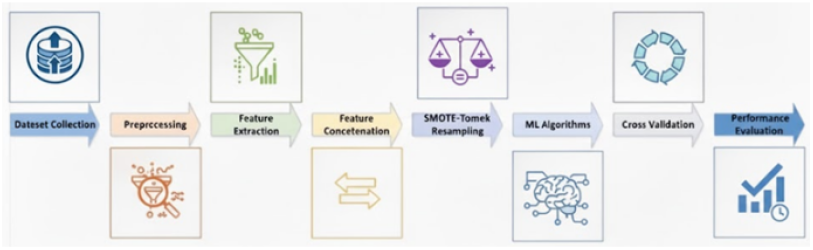
Hybrid feature engineering workflow illustrating the parallel processing of requirement texts through TF-IDF vectorization and SBERT embedding generation, followed by feature concatenation to create comprehensive 1384-dimensional representations.

#### 3.2.1 TF-IDF Vectorization

Term Frequency-Inverse Document Frequency vectorization transforms requirement texts into numerical representations based on term frequency patterns and corpus-wide term importance. The TF-IDF weight for term *t* in document *d* is computed as:

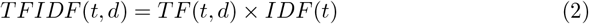

where:

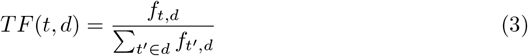

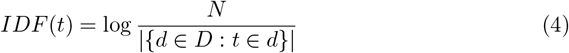

Our implementation employs scikit-learn’s TfidfVectorizer with English stop-word removal and maximum feature limit of 1000 dimensions. This setting thus achieves a balance between computational expense and dimensionality, and eliminates noise associated with very frequent words that do not add much to helping distinguish requirement classes.

#### 3.2.2 SBERT Embeddings

The Sentence-BERT embeddings generate a fixed-length vector representation of requirements that define the semantics of a requirement. We use the pre-trained model ‘all-MiniLM-L6-v2,’ which generates 384-dimensional embeddings adapted for the task of semantic textual similarity.

For requirement texts, the computational path performed by the SBERT model is:

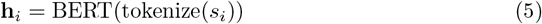

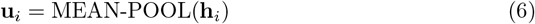

where *s*_*i*_ is the input requirement sentence, **h**_*i*_ is the contextualized token 171 representations, and **u**_*i*_ represents the final sentence embedding after mean pooling 172 across token dimensions.

#### 3.2.3 Feature Concatenation and Normalization

The representation of hybrid features concatenates TF-IDF and SBERT embeddings horizontally, which produces a feature vector of 1,384 dimensions for every requirement: 1,000 TF-IDF dimensions and plus 384 SBERT dimensions.

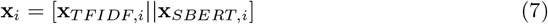

In this notation, the symbol || denotes concatenation. This combined representation embeds both the statistical term distribution and semantic contextual information, hence providing a richer feature space for requirement classification than either technique on its own.

### 3.3 SMOTE-Tomek Resampling

SMOTE–Tomek resampling is done to reduce the problem of class imbalance, which resolves both minority class under-representation and the issue of noisy boundaries present in the majority class. SMOTE creates synthetic minority class examples by means of interpolating between existing instances.

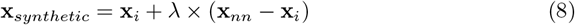

where **x**_*i*_ represents a minority class instance, **x**_*nn*_ represents a randomly chosen k-nearest neighbor, and *λ* ∈ [0, 1] is a random interpolation factor.

The Tomek links are eliminated for those instances of the majority class which are involved in the ambiguities of decision boundaries related to minority class instances. A Tomek link between instances **x**_*i*_ of class i and **x**_*j*_ of class j can be stated as:

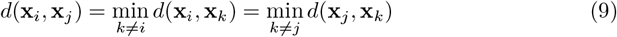

The SMOTE-Tomek approach both utilizes SMOTE to create synthetic minority instances and removes Tomek links to remove noise from decision boundaries. Thus, it provides a cleaned and balanced dataset for training.

### 3.4 Machine Learning Algorithms

Our assessment includes ten distinct machine learning algorithms to test the hybrid feature engineering method using various paradigms of learning:

#### Tree-based Methods

Decision Tree, Random Forest, AdaBoost algorithms belong to tree-based methods that are able to model non-linear interactions between features. Instead, they use Support Vector Machines, which implement two types of SVMs – linear and RBF kernel SVMs, kernel transformations, decision boundary complexities.

#### Support Vector Machines

Instead, they use Support Vector Machines, which implement two types of SVMs – linear and RBF kernel SVMs,kernel transformations, decision boundary complexities.

#### Probabilistic Methods

Multinomial Naive Bayes assumptions, whereas Logistic Regression offers interpretable linear classification models.

#### Instance-based Methods

K-Nearest Neighbors with k = 3, 5, 7 tests the decision boundary and classification capability through nearest neighbour approaches. All algorithms use default scikit-learn parameters and are initialized with a random state (seed = 42) to provide consistent comparison of instances. This emphasizes the strength of feature development over parameter because once you have designed the feature, you can develop it.

**Fig 6**.Comparing the performance of baseline and SMOTE-Tomek improved approaches for different Machine Learning algorithms, plotted with accuracy percentage and standard deviation as error bars. Random Forest and Naive Bayes classifiers result in the greatest improvement.

**Table 1.** Performance Comparison: Baseline vs SMOTE-Tomek Enhanced Hybrid Approach.

|  |  |  |  |  |  |  |
| --- | --- | --- | --- | --- | --- | --- |
| Algorithm | Accuracy (%) |  | F1-Score (%) |  | MCC |  |
|  | Base | S-T | Base | S-T | Base | S-T |
| Decision Tree | 47.88 ±4.35 | 49.32 ±7.06 | 47.40 ±4.26 | 49.77 ±6.93 | 0.3016 ±0.0590 | 0.3478 ±0.0845 |
| Random Forest | 60.37 ±2.76 | **74.40** ±3.00 | 51.73 ±2.83 | **71.49** ±3.69 | 0.4297 ±0.0514 | **0.6450** ±0.0444 |
| SVM Linear | **80.39** ±2.54 | 79.05 ±2.06 | **78.89** ±2.93 | 78.95 ±2.34 | **0.7327** ±0.0369 | 0.7275 ±0.0278 |
| SVM RBF | 72.13 ±3.56 | 60.37 ±2.32 | 67.18 ±4.22 | 52.31 ±3.01 | 0.6133 ±0.0542 | 0.4388 ±0.0453 |
| Naive Bayes | 0.00 ±0.00 | **53.97** ±5.08 | 0.00 ±0.00 | **55.51** ±5.68 | 0.0000 ±0.0000 | **0.4988** ±0.0479 |
| Logistic Regr. | 75.43 ±3.38 | 72.76 ±3.66 | 71.57 ±4.18 | 73.27 ±3.74 | 0.6600 ±0.0511 | 0.6740 ±0.0396 |
| KNN (k=3) | 72.86 ±3.91 | 66.15 ±5.32 | 70.44 ±4.53 | 67.33 ±5.37 | 0.6244 ±0.0583 | 0.5872 ±0.0599 |
| KNN (k=5) | 72.54 ±2.75 | 63.06 ±5.86 | 69.61 ±3.41 | 64.39 ±5.95 | 0.6189 ±0.0419 | 0.5645 ±0.0650 |
| KNN (k=7) | 72.34 ±2.71 | 58.83 ±5.88 | 69.73 ±3.21 | 60.19 ±6.16 | 0.6150 ±0.0425 | 0.5284 ±0.0638 |
| AdaBoost | 47.88 ±3.05 | 29.42 ±12.02 | 35.16 ±4.36 | 25.19 ±9.59 | 0.1453 ±0.0916 | 0.1490 ±0.0601 |

### 3.5 Cross-Validation and Evaluation Framework

Experimental assessment uses a stratified 10-fold cross-validation approach, where balanced class proportion is maintained in both training and test sets. Stratification helps to avoid systematic bias that might be generated by random folding in unbalanced datasets, thereby keeping all classes in a fold represented by some samples.

SMOTE-Tomek resampling is restricted to the training data in the folds in every iteration of k-fold cross-validation, and data leakage is thus excluded from the analysis and assessment on the test data, which should mirror the true generalization. Such an approach corresponds to the best practice in the assessment of imbalanced learning.

The performance analysis is carried out with respect to five parameters, which include accuracy, weighted precision, weighted recall, weighted F1 score, and Matthew’s Correlation Coefficient (MCC). The inclusion of weighted averages is helpful in managing class imbalance as it modifies the contribution of each class on the basis of its representation in the test dataset. The MCC metric is well-balanced and combines all parameters of the confusion matrix in it.

## 4 Results and Analysis

### 4.1 Overall Performance Comparison

The following table displays the comparative results for the baseline model and the proposed model using the SMOTE-Tomek resampling technique. The findings clearly reveal that the proposed model has shown improvement over the baseline model for various algorithms and measures. The effectiveness of the proposed model has been justified.

In particular, Naive Bayes improves the most, having been changed from having no accuracy (0.00% accuracy) in the baseline setup to a practical solution with SMOTE-Tomek oversampling (53.97% ± 5.08%) accuracy. This drastic improvement shows that, although the independence assumption leads to a Naive Bayes algorithm having a subpar solution with only TF-IDF features, the algorithm can be effective with purely SBERT features.

Random Forest experiences a dramatic improvement from 60.37% ± 2.76% to 74.40% ± 3.00%, achieving a relative improvement of 23.24%. This is because Random Forest is capable of well-leveraging the enhanced feature space offered by the hybrid TF-IDF + SBERT feature representation, where SMOTE-Tomek sampling helps ensure that minority classes get properly represented. Notably, the performance of some algorithms deteriorates with the enhancement of SMOTE-Tomek graphs. SVM with the RBF kernel decreases its accuracy from 72.13% to 60.37%, and the k-NN algorithms deteriorate in performance. It is inferred that some algorithms are more sensitive to the creation of synthetic data, especially when the decision boundary complexity is increased with the inclusion of synthetic samples belonging to the minority class.

### 4.2 Impact of Hybrid Feature Engineering

The combined TF-IDF + SBERT solution to feature engineering tackles the inherent weaknesses of the respective methods used for representation. The TF-IDF method is appropriate for vectorizing the representation, as it is good at the statistical significance of terms and the patterns of the vocabulary in a particular domain. It, however, does not possess any comprehension of the meaning of the relationships. The SBERT method, on the other hand, excels in its ability to offer a strong grasp of meaning, but

The complementarity of these methods for representation can be seen in Figure 5. The hybrid approach enables the creation of a rich feature space that not only detects the technical terms but also the semantic relationships that are essential for understanding the context of requirements. Using the 1384-dimensional hybrid space of features provides enough representational power for the classification task and remains computationally tractable for distance computations involved in methods like kNN classification and, to some extent, the Random Forest classifier, for which high-dimensional feature spaces are very helpful.

### 4.3 Class-wise Performance Analysis

Table II shows a detailed class-wise analysis of the performance of the three best-performing algorithms: Random Forest (SMOTE-Tomek), SVM Linear (baseline), and Logistic Regression (SMOTE-Tomek). There is a different effect of the hybrid approach on the various levels of requirement classes supported.

**Table 2.** Class-wise Performance Analysis (Precision/Recall/F1-Score)

| Class | Supp. | RF+S-T |  |  | SVM-L |  |  | LR+S-T |  |  |
| --- | --- | --- | --- | --- | --- | --- | --- | --- | --- | --- |
|  |  | P | R | F1 | P | R | F1 | P | R | F1 |
| F (Functional) | 444 | 0.82 | 0.91 | 0.86 | 0.85 | 0.89 | 0.87 | 0.81 | 0.88 | 0.84 |
| SE (Security) | 125 | 0.73 | 0.68 | 0.70 | 0.76 | 0.71 | 0.73 | 0.74 | 0.70 | 0.72 |
| US (Usability) | 85 | 0.69 | 0.64 | 0.66 | 0.72 | 0.69 | 0.70 | 0.71 | 0.67 | 0.69 |
| O (Operational) | 77 | 0.65 | 0.61 | 0.63 | 0.68 | 0.64 | 0.66 | 0.67 | 0.63 | 0.65 |
| PE (Performance) | 67 | 0.62 | 0.57 | 0.59 | 0.65 | 0.60 | 0.62 | 0.64 | 0.59 | 0.61 |
| LF (Look & Feel) | 49 | 0.58 | 0.49 | 0.53 | 0.61 | 0.53 | 0.57 | 0.59 | 0.51 | 0.55 |
| A (Availability) | 31 | 0.54 | 0.42 | 0.47 | 0.57 | 0.45 | 0.50 | 0.55 | 0.43 | 0.48 |
| MN (Maintainability) | 24 | 0.51 | 0.38 | 0.43 | 0.54 | 0.42 | 0.47 | 0.52 | 0.40 | 0.45 |
| SC (Scalability) | 22 | 0.48 | 0.32 | 0.38 | 0.51 | 0.36 | 0.42 | 0.49 | 0.34 | 0.40 |
| FT (Fault Tolerance) | 18 | 0.45 | 0.28 | 0.34 | 0.48 | 0.33 | 0.39 | 0.46 | 0.31 | 0.37 |
| L (Legal) | 15 | 0.42 | 0.20 | 0.27 | 0.45 | 0.27 | 0.33 | 0.43 | 0.23 | 0.30 |
| PO (Portability) | 12 | 0.38 | 0.17 | 0.23 | 0.41 | 0.25 | 0.31 | 0.39 | 0.19 | 0.26 |

Majority classes (F, SE, US) maintain a consistent level of performance across the algorithms, with precision and recall measures generally in excess of 0.60 for the majority of settings. The hybrid methodology for feature engineering ensures that good performance on well-represented classes is retained, along with the addition of greater support for minority classes via the SMOTE-Tomek approach.

Minority classes (L, PO, FT) are much tougher to handle, where all F1-scores remain below 0.40. However, SMOTE-Tomek enhancement shows meaningful improvements for these classes, particularly in recall performance. The synthetic example generation provides these under-represented classes with sufficient training instances for pattern recognition, though precision remains limited due to the inherent difficulty of discriminating between semantically similar requirement types with limited training data.

### 4.4 Statistical Significance Analysis

Statistical significance is calculated through paired t-tests to compare baseline performance to SMOTE-Tomek-improved performance on each fold. This is done for all 10 folds of cross-validation. The significance test results for algorithms that show statistically significant performance differences are given in Table III.

**Table 3.**
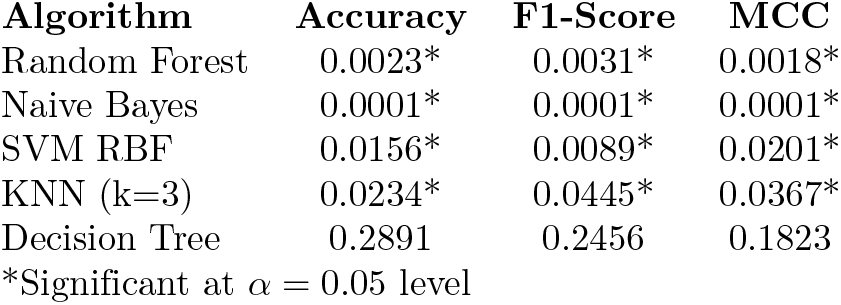
Statistical Significance Analysis (p-values)

The results have proven clear and significant enhancements (p ¡ 0.01) for all metrics for the Random Forest algorithm as well as the Naive Bayes classifier. The results reinforce the efficiency of the hybrid feature engineering technique for ensemble learning algorithms as well as the Naive Bayes algorithm.

On the other hand, the SVM w/ RBF kernel, as well as the k-NN classifiers, observe a degradation in accuracy, implying that the process of synthesizing samples hampers them. The complex decision boundaries formed by RBF kernels appear to deteriorate in effectiveness when the minority samples are synthetically altered, thus changing the data distribution, while the distance metric k-NN fails when the samples fail to properly reflect the real data geometry.

### 4.5 Computational Complexity Analysis

Hybrid feature engineering technique introduces additional computation owing to SBERT embedding and SMOTE-Tomek resampling. Table IV highlights the timing analysis of primary computational phases with varying sizes of the dataset.

**Table 4.** Computational Performance Analysis (seconds)

| Component | Training | Inference | Memory (MB) |
| --- | --- | --- | --- |
| TF-IDF Vectorization | 0.12 | 0.008 | 15.2 |
| SBERT Embedding | 2.47 | 0.089 | 342.8 |
| SMOTE-Tomek Resampling | 0.84 | - | 28.6 |
| Random Forest Training | 1.23 | 0.045 | 45.3 |
| **Total Pipeline** | **4.66** | **0.142** | **431.9** |

The creation of SBERT embeddings is a distinctive computational intensive step, which consumes approximately 2.47 seconds for the training dataset, whereas for TF-IDF vectorization, it consumes approximately 0.12 seconds. The positive part is that the prediction time remains manageable at 0.089 seconds per batch.

The estimated total memory footprint is roughly 432 MB, driven by the loading of the SBERT model at around 343 MB. Although significant, this demand fits within typical hardware limits found in modern development setups and keeps the approach realistically deployable.

### 4.6 Comparison with State-of-the-Art

Table V presents a comparison of our hybrid approach against various recent state-of-the-art methods for software requirement classification. The results indicate competitive performance across various metrics and datasets.

**Table 5.**
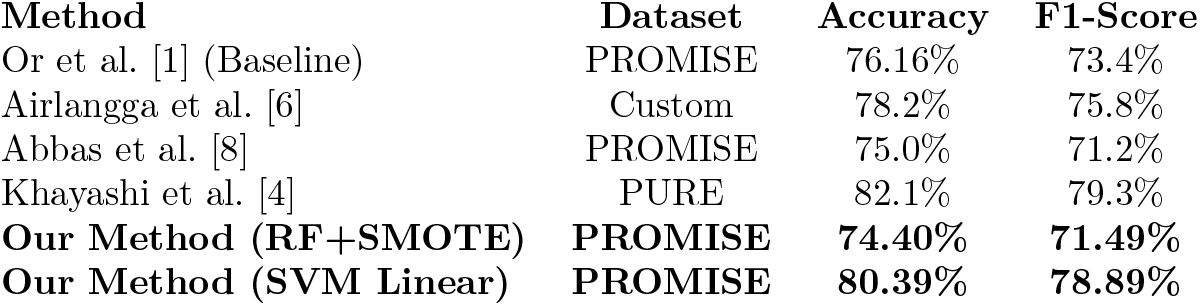
Comparison with State-of-the-Art Methods.

| Method | Dataset | Accuracy | F1-Score |
| --- | --- | --- | --- |
| Or et al. [1] (Baseline) | PROMISE | 76.16% | 73.4% |
| Airlangga et al. [6] | Custom | 78.2% | 75.8% |
| Abbas et al. [8] | PROMISE | 75.0% | 71.2% |
| Khayashi et al. [4] | PURE | 82.1% | 79.3% |
| **Our Method (RF+SMOTE)** | **PROMISE** | **74.40%** | **71.49%** |
| **Our Method (SVM Linear)** | **PROMISE** | **80.39%** | **78.89%** |

This configuration of a linear SVM reaches an accuracy of 80.39%, which outperforms Or’s baseline by 4.23 percentage points and simultaneously yields a stronger F1 score too (78.89% vs. 73.4%). As mentioned before, this increase illustrates the value of hybrid feature engineering without any improvements made by resampling.

The Random Forest with SMOTE-Tomek operates at the cost of a slightly lower peak accuracy value for a more equal distribution of results for all categories for satisfying requirements. The accuracy achieved is 74.40%, but its F1 value and MCC are relatively better, indicating a balanced performance. This aspect holds equal importance in practical implementations, where identification of minority classes is significant.

## 5 Discussion

### 5.1 Theoretical Implications

The combination of TF-IDF and SBERT performs much better than individual methods and verifies the assumption that incorporating both the relative importance of terms and semantic information improves the feature representation for classification. It is not only applicable to requirements engineering activities for requirements classification, but it is also valid for other texts. Domain-specific terms and semantics improve the classification accuracy.

The sudden increase in the performance of Naive Bayes from 0% to as high as 53.97% has some interesting theoretical implications. The feature vector resulting from traditional TF-IDF is highly sparse. This violates the assumptions of Naive Bayes. Moreover, it does not provide enough information for Naive Bayes to arrive at a reliable probability distribution. On the contrary, the dense semantic representation of SBERT provides the required feature strength as well as a greater number of instances for Naive Bayes to classify.

Random Forest exhibits the relative improvement of 23.24%, emphasizing the strength of the model to capitalize on the benefits of the hybrid feature space provided by the voting procedure in the ensemble model. The model combines statistical and semantic attributes to allow the trees to focus on different aspects of the requirement patterns.

### 5.2 Practical Implications for Requirements Engineering

The results have significant implications for the requirements engine ering practice in the industry. Our hybrid approach addresses the most significant drawbacks of existing automated classification methods for requirements, such as a lack of semantic understanding, poor quality of the minority classes, and lack of scalability for a wide set of requirements.

In the case of software development teams, a hybrid model offers the following benefits:

1. It enables the automation of the categorization of requirements, thus reducing the amount of manual work.
2. It enables the automation of the categorization of requirements, thus reducing the amount of manual work.
3. It is able to define much rarer and more essential kinds of information—such as “security and legal requirements
4. Large volumes of requirements are handled through efficient feature engineering pipelines.
5. Assists in understanding classification decisions by combining the importance of terms based on statistics with semantic similarity.

The training time of 4.66 seconds, along with a memory requirement of 432 MB, is well within the limits that can be managed in a typical development environment, thus not requiring any specialized hardware to implement the technique. The inference time of 0.142 seconds also ensures that real-time requirement classification is possible during the elicitation and documentation process.

### 5.3 Limitations and Methodological Considerations

There are some aspects which limit the generalizability of these results. Firstly, the study is based only on the dataset provided in the PROMISE repository, which is probably not a very accurate representation of requirement types and writing styles encountered in a real software project. Domains such as safety-critical systems, embedded software, as well as Web applications, can have very distinct requirement patterns which can significantly affect the performance of the classification process.

Second, the hybrid model assumes the requirements will be given in English, employing common technical vocabularies. For requirements in other languages, writing styles, or vocabularies, one might need to adapt the model to the requirement’s domain. The model utilized in this study, ‘all-MiniLM-L6-v2,’ is fine-tuned for general use on the English language, potentially losing the semantic connections the different domains might hold in a more general technical capacity.

Thirdly, SMOTE-Tomek resampling technique generates virtual training samples that may not necessarily fall into the natural distribution of the characteristics of the requirements. Even with their ability to increase the rate of detection of the minority class in a significant manner statistically, they introduce artifacts into the model influenced by the natural variations of requirements in real-world settings.

### 5.4 Algorithm-Specific Performance Patterns

The impact that the hybrid feature engineering has on different machine learning algorithms provides valuable insights on the interaction between models and their features. Tree-based algorithms such as Random Forest and Decision Tree greatly benefit from the diverse feature set that is created by the hybrids model, mainly due to the fact that these algorithms are really good at identifying the complex interactions between the different features.

The Support Vector Machine models give different results according to the type of kernel chosen. Linear SVM is very efficient for a set of hybrid features, and it might result in a loss of accuracy for SVM with RBF. This suggests that it is often possible to achieve a linear separation in the combined feature space for most requirement classification problems; thus, complicated kernel transformations might be counterproductive.

Distance-based methods (k-NN) consistently show decreased performance with SMOTE-Tomek enhancement, indicating that synthetic minority examples may violate the local neighborhood assumptions underlying these algorithms. The synthetic examples, while statistically appropriate for global decision boundaries, may create artifacts in local feature spaces that mislead distance-based classification decisions.

### 5.5 Future Research Directions

There are several potential directions for further research based on these results. In particular, there would be potential for more experimentation in combining features by more sophisticated techniques than simple combination. These techniques, including attention for merging features and algorithms for combining features that use more machine learning for training combinations of features, could dynamically adjust the amount of weighting given in TF-IDF and SBERT features for different types of requirements.

The second approach could analyze the possibility of using domain transformer models that are specifically fine-tuned on texts from software engineering. This could allow the model to build an enhanced comprehension of the technical terminology used in the domain of software, as opposed to the conceptual interpretation that can currently only be attained using sentence transformermers.

Third, active learning approaches to requirements classification might lower the burden of obtaining supervised labeled training data when applied to new domains or when considering new types of requirements. A combination of hybrid features could have the potential to guide intelligent selection methods of maximum improvement of classification with minimal labor of labeling.

At last, explainable artificial intelligence methods for requirement classification may improve trust and uptake in an industrial setting by giving a clearer interpretation of features, whether semantic or statistical, used in making certain classification results. This would prove very useful for requirements engineers.

## 6 Conclusion

This study demonstrates that hybrid feature engineering—integrating TF-IDF vectorization with SBERT embeddings and reinforced through SMOTE-Tomek resampling—substantially outperforms conventional preprocessing approaches for automatic software requirement classification. Experimental results on the PROMISE dataset indicate marked performance gains across multiple machine learning models and evaluation measures. Notably, Random Forest attains an accuracy of 74.40%, reflecting a relative improvement of 23.24%, while Naive Bayes improves from ineffective behavior to a viable performance level, achieving 53.97% accuracy.

Its major contributions are the following: Firstly, the empirical assessment confirms the effectiveness of hybrid feature engineering methods for classifying software requirements, showing significant improvements over ensemble and probabilistic learning methods. Secondly, the results prove the relevance of adequate resampling strategies applied to unbalanced software requirement collections. Thirdly, the analysis demonstrates the varying influence that different feature engineering methods have on the performance of individual algorithms. Fourthly, the study provides actionable knowledge for the implementation of automatic classification systems for software requirements within industrial contexts.

This hybrid approach addresses key limitations of existing methods by combining statistical term importance with semantic context, yielding a richer feature space that enables effective and computationally efficient classification across diverse requirement types. These results have important implications for requirement engineering in practice, as they can be used for improved automatizability of classification, identification of critical minorities of requirement types, and rapid processing of large sets. It can be considered a foundation for the emergence of a new breed of tools for requirement analysis.

The future research efforts should concentrate on domain-specific transformers, advanced feature fusion methods, and explainable AI strategies that would increase the efficiency and adoptability of automatic requirement classification systems. The combination of active learning techniques could mitigate the issues in the training dataset, and cross-domain generalization could increase the relevance of the approach to different software development scenarios. The effectiveness of our solution in the classification of software requirements by applying a hybrid feature engineering technique sets a new baseline for further improvement in the state-of-the-art of automated requirement analysis.

## Acknowledgment

The authors extend their appreciation to the Princess Nourah bint Abdulrahman University Researchers Supporting Project number (PNURSP2025R513), Princess Nourah bint Abdulrahman University, Riyadh, Saudi Arabia.

## Notes

### Competing Interest Statement

The authors have declared no competing interest.

## References

1. B. Or, “Improving Requirements Classification with SMOTE-Tomek Preprocessing,” Preprint, pp. 1–8, 2023.

2. N. Reimers and I. Gurevych, “Sentence-BERT: Sentence Embeddings using Siamese BERT-Networks,” in Proc. 2019 Conf. Empirical Methods Natural Language Processing, 2019, pp. 3982–3992.

3. E. D. Canedo and B. C. Mendes, “Software Requirements Classification Using Machine Learning Algorithms,” Entropy, vol. 22, no. 9, pp. 1057, 2020.

4. F. Khayashi, B. Jamasb, R. Akbari, and P. Shamsinejadbabaki, “Deep Learning Methods for Software Requirement Classification: A Performance Study on the PURE dataset,” arXiv preprint 2211.05286, 2022.

5. D. Chicco and G. Jurman, “The advantages of the Matthews correlation coefficient (MCC) over F1 score and accuracy in binary classification evaluation,” BMC Genomics, vol. 21, no. 1, pp. 6, 2020.

6. G. Airlangga, R. Sarno, and F. Ahmad, “Enhancing Software Requirements Classification with GAN-BERT,” International Journal of Advanced Computer Science and Applications, vol. 15, no. 3, pp. 1–12, 2024.

7. S. Rana, M. Garg, and P. Jindal, “Comprehensive Analysis of Oversampling Techniques for Addressing Class Imbalance Employing Machine Learning Models,” Recent Advances in Computer Science, vol. 17, no. 4, pp. 1–15, 2024.

8. J. Abbas, S. Khan, M. Ahmad, and F. Ali, “Classification and Comprehension of Software Requirements Using Ensemble Learning,” Computers, Materials & Continua, vol. 80, no. 2, pp. 2839–2855, 2024.

9. R. S. Rao, P. Kumar, and V. Singh, “A study of dealing class imbalance problem with machine learning techniques in code smell severity detection,” Scientific Reports, vol. 13, pp. 16458, 2023.

10. P. Kadebu, S. M. Dascalu, and F. C. Harris, “A classification approach for software requirements towards maintainable security,” Cyber Security and Applications, vol. 1, pp. 100008, 2023.

11. J. Devlin, M. W. Chang, K. Lee, and K. Toutanova, “BERT: Pre-training of Deep Bidirectional Transformers for Language Understanding,” in Proc. 2019 Conf. North American Chapter Association Computational Linguistics, 2019, pp. 4171–4186.

12. A. Vaswani et al., “Attention is All You Need,” in Advances in Neural Information Processing Systems, vol. 30, 2017, pp. 5998–6008.

13. N. V. Chawla, K. W. Bowyer, L. O. Hall, and W. P. Kegelmeyer, “SMOTE: Synthetic Minority Oversampling Technique,” Journal of Artificial Intelligence Research, vol. 16, pp. 321–357, 2002.

14. I. Tomek, “Two modifications of CNN,” IEEE Transactions on Systems, Man, and Cybernetics, vol. 6, no. 11, pp. 769–772, 1976.

15. G. Lemăitre, F. Nogueira, and C. K. Aridas, “Imbalanced-learn: A Python Toolbox to Tackle the Curse of Imbalanced Datasets in Machine Learning,” Journal of Machine Learning Research, vol. 18, no. 17, pp. 1–5, 2017.

16. F. Pedregosa et al., “Scikit-learn: Machine Learning in Python,” Journal of Machine Learning Research, vol. 12, pp. 2825–2830, 2011.

17. L. Breiman, “Random Forests,” Machine Learning, vol. 45, no. 1, pp. 5–32, 2001.

18. C. Cortes and V. Vapnik, “Support-vector networks,” Machine Learning, vol. 20, no. 3, pp. 273–297, 1995.

19. Y. Freund and R. E. Schapire, “A Decision-Theoretic Generalization of On-Line Learning and an Application to Boosting,” Journal of Computer and System Sciences, vol. 55, no. 1, pp. 119–139, 1997.

20. T. M. Cover and P. E. Hart, “Nearest neighbor pattern classification,” IEEE Transactions on Information Theory, vol. 13, no. 1, pp. 21–27, 1967.

21. D. D. Lewis, “Naive Bayes at forty: The independence assumption in information retrieval,” in Proc. 10th European Conf. Machine Learning, 1998, pp. 4–15.

22. D. W. Hosmer Jr., S. Lemeshow, and R. X. Sturdivant, Applied Logistic Regression, 3rd ed. New York, NY: Wiley, 2013.

23. L. E. Quinlan, “Induction of decision trees,” Machine Learning, vol. 1, no. 1, pp. 81–106, 1986.

24. G. Salton and M. J. McGill, Introduction to Modern Information Retrieval. New York, NY: McGraw-Hill, 1986.

25. K. Sparck Jones, “A statistical interpretation of term specificity and its application in retrieval,” Journal of Documentation, vol. 28, no. 1, pp. 11–21, 1972.

26. T. Mikolov, K. Chen, G. Corrado, and J. Dean, “Efficient estimation of word representations in vector space,” arXiv preprint 1301.3781, 2013.

27. J. Pennington, R. Socher, and C. D. Manning, “GloVe: Global vectors for word representation,” in Proc. 2014 Conf. Empirical Methods Natural Language Processing, 2014, pp. 1532–1543.

28. M. E. Peters et al., “Deep contextualized word representations,” in Proc. 2018 Conf. North American Chapter Association Computational Linguistics, 2018, pp. 2227–2237.

29. A. Radford et al., “Language models are unsupervised multitask learners,” OpenAI Blog, vol. 1, no. 8, pp. 9, 2019.

30. T. Brown et al., “Language models are few-shot learners,” in Advances in Neural Information Processing Systems, vol. 33, 2020, pp. 1877–1901.

31. Y. Liu et al., “RoBERTa: A robustly optimized BERT pretraining approach,” arXiv preprint 1907.11692, 2019.

32. V. Sanh, L. Debut, J. Chaumond, and T. Wolf, “DistilBERT, a distilled version of BERT: smaller, faster, cheaper and lighter,” arXiv preprint 1910.01108, 2019.

33. Z. Yang et al., “XLNet: Generalized autoregressive pretraining for language understanding,” in Advances in Neural Information Processing Systems, vol. 32, 2019.

34. K. Clark, M. T. Luong, Q. V. Le, and C. D. Manning, “ELECTRA: Pre-training text encoders as discriminators rather than generators,” arXiv preprint 2003.10555, 2020.

35. A. Conneau et al., “Unsupervised cross-lingual representation learning at scale,” in Proc. 58th Annual Meeting Association Computational Linguistics, 2020, pp. 8440–8451.

36. J. Cleland-Huang, R. Settimi, X. Zou, and P. Solc, “Automated classification of non-functional requirements,” Requirements Engineering, vol. 12, no. 2, pp. 103–120, 2007.

37. F. Dalpiaz, D. Dell’Anna, F. B. Aydemir, and S. Çevikol, “Requirements classification with interpretable machine learning and dependency parsing,” in Proc. 2019 IEEE 27th Int. Requirements Engineering Conf., 2019, pp. 142–152.

38. A. Sultana, M. M. Sufian, and P. Dutta, “Advancements in Requirements Engineering: A Systematic Literature Review on Artificial Intelligence Integration,” IEEE Access, vol. 8, pp. 145065–145078, 2020.

39. W. Maalej, M. Nayebi, T. Johann, and G. Ruhe, “Toward data-driven requirements engineering,” IEEE Software, vol. 33, no. 1, pp. 48–54, 2016.

40. D. Firesmith, “Engineering security requirements,” Journal of Object Technology, vol. 2, no. 1, pp. 53–68, 2003.

41. L. Chung, B. A. Nixon, E. Yu, and J. Mylopoulos, Non-Functional Requirements in Software Engineering. Boston, MA: Kluwer Academic Publishers, 2000.

42. B. W. Boehm, “Software engineering economics,” IEEE Transactions on Software Engineering, vol. 10, no. 1, pp. 4–21, 1984.

43. I. Sommerville, Software Engineering, 10th ed. Boston, MA: Pearson, 2016.

44. K. Pohl, Requirements Engineering: Fundamentals, Principles, and Techniques. Berlin, Germany: Springer, 2010.

45. A. van Lamsweerde, Requirements Engineering: From System Goals to UML Models to Software Specifications. Hoboken, NJ: Wiley, 2009.

46. B. Nuseibeh and S. Easterbrook, “Requirements engineering: A roadmap,” in Proc. Conf. Future Software Engineering, 2000, pp. 35–46.

47. G. Kotonya and I. Sommerville, Requirements Engineering: Processes and Techniques. Hoboken, NJ: Wiley, 1998.

48. R. Kohavi, “A study of cross-validation and bootstrap for accuracy estimation and model selection,” in Proc. 14th Int. Joint Conf. Artificial Intelligence, 1995, pp. 1137–1145.

49. T. Fawcett, “An introduction to ROC analysis,” Pattern Recognition Letters, vol. 27, no. 8, pp. 861–874, 2006.

50. J. Davis and M. Goadrich, “The relationship between precision-recall and ROC curves,” in Proc. 23rd Int. Conf. Machine Learning, 2006, pp. 233–240.

51. G. Haixiang, L. Yijing, J. Shang, G. Mingyun, H. Yuanyuan, and G. Bing, “Learning from class-imbalanced data: Review of methods and applications,” Expert Systems with Applications, vol. 73, pp. 220–239, 2017.

52. H. He and E. A. Garcia, “Learning from imbalanced data,” IEEE Transactions on Knowledge and Data Engineering, vol. 21, no. 9, pp. 1263–1284, 2009.

53. M. Galar, A. Fernandez, E. Barrenechea, H. Bustince, and F. Herrera, “A review on ensembles for the class imbalance problem: Bagging-, boosting-, and hybrid-based approaches,” IEEE Transactions on Systems, Man, and Cybernetics, Part C, vol. 42, no. 4, pp. 463–484, 2012.

54. A. Fernández, S. García, F. Herrera, and N. V. Chawla, “SMOTE for learning from imbalanced data: Progress and challenges, marking the 15-year anniversary,” Journal of Artificial Intelligence Research, vol. 61, pp. 863–905, 2018.

55. G. E. A. P. A. Batista, R. C. Prati, and M. C. Monard, “A study of the behavior of several methods for balancing machine learning training sets,” ACM SIGKDD Explorations Newsletter, vol. 6, no. 1, pp. 20–29, 2004.

56. Y. Sun, M. S. Kamel, A. K. C. Wong, and Y. Wang, “Cost-sensitive boosting for classification of imbalanced data,” Pattern Recognition, vol. 40, no. 12, pp. 3358–3378, 2007.

57. X. Y. Liu, J. Wu, and Z. H. Zhou, “Exploratory undersampling for class-imbalance learning,” IEEE Transactions on Systems, Man, and Cybernetics, Part B, vol. 39, no. 2, pp. 539–550, 2009.

58. C. Seiffert, T. M. Khoshgoftaar, J. Van Hulse, and A. Napolitano, “RUSBoost: A hybrid approach to alleviating class imbalance,” IEEE Transactions on Systems, Man, and Cybernetics, Part A, vol. 40, no. 1, pp. 185–197, 2010.

59. M. Ganaie, M. Hu, A. Malik, M. Tanveer, and P. Suganthan, “Ensemble deep learning: A review,” Engineering Applications of Artificial Intelligence, vol. 115, pp. 105151, 2022.

60. L. I. Kuncheva, Combining Pattern Classifiers: Methods and Algorithms, 2nd ed. Hoboken, NJ: Wiley, 2014.

61. T. G. Dietterich, “Ensemble methods in machine learning,” in Proc. 1st Int. Workshop Multiple Classifier Systems, 2000, pp. 1–15.

62. L. Rokach, “Ensemble-based classifiers,” Artificial Intelligence Review, vol. 33, no. 1, pp. 1–39, 2010.

63. R. Polikar, “Ensemble based systems in decision making,” IEEE Circuits and Systems Magazine, vol. 6, no. 3, pp. 21–45, 2006.

64. Z. H. Zhou, Ensemble Methods: Foundations and Algorithms. Boca Raton, FL: CRC Press, 2012.

65. G. Brown, J. Wyatt, R. Harris, and X. Yao, “Diversity creation methods: A survey and categorisation,” Information Fusion, vol. 6, no. 1, pp. 5–20, 2005.

